# Competitive HIV budding suggests that a self-packaging gRNA:Gag-Pol complex directs HIV assembly and enforces infectivity

**DOI:** 10.1101/2022.08.03.502595

**Authors:** Haley Durden, Ipsita Saha, Benjamin Preece, Brian MacArthur, Abby Petersen, Wiley Peppel, Rodrigo Gallegos, Saveez Saffarian

**Affiliations:** Center for Cell and Genome Science, University of Utah, Salt Lake City, UT, 84112, USA; Department of Physics and Astronomy, University of Utah, Salt Lake City, UT, 84112, USA; Laboratory of Cell and Developmental Signaling, Center for Cancer Research, National Cancer Institute, National Institutes of Health, Frederick, MD, 21702, USA; School of Biological Sciences, University of Utah, Salt Lake City, UT, 84112, USA

## Abstract

To resolve the assembly mechanism of infectious HIV virions, we tested the ability of HIV to assemble infectious virions in the presence of a titrated mix of infectious/ non-infectious proviral genomes. The analysis of our assembly competitions shows that during translation, 15 ± 5 Gag-Pols bind back to their parental gRNA creating a gRNA_1_: Gag-Pol_15_ complex. This complex initiates the infectious virion assembly through interactions mediated by cis packaged Gag/Gag-pols and the plasma membrane. Our analysis also shows the number of Gag-Pol and Env proteins packaged in an infectious HIV virion and the minimum functional units of these proteins required for viral infectivity. We suggest that aside from orchestrating the infectious virion assembly the gRNA_1_: Gag-Pol_15_ complex plays a major role in HIV evolution and likely hampers effectiveness of antiviral therapies.

**One-Sentence Summary:** Competitive HIV budding reveals gRNA_1_: Gag-Pol_15_ complex orchestrating viral assembly.

## Introduction

Acute Immune Deficiency Syndrome (AIDS) is caused by Human immunodeficiency virus (HIV) which continues to be a major pathogen with a significant death toll around the globe (WHO, 2021). Currently, inhibiting key HIV interactions serves as the basis of antiviral therapies for AIDS (Campbell and Hope, 2015; Kleinpeter and Freed, 2020). Directly observing HIV’s protein-protein, protein-lipid and protein-RNA interactions within the context of fully infectious HIV replication is challenging, mainly because wild type HIV loses significant infectivity when its gRNA or viral proteins are tagged (Bendjennat and Saffarian, 2019; Hubner et al., 2009; Marsden et al., 2020; McDonald et al., 2002; Müller et al., 2004). The current model of HIV budding, therefore, envisions HIV budding as a sum of independently verified interactions (Freed, 2015). Primed by the observed resistance of HIV gRNA to modifications, we asked if there are cooperative and long-range molecular interactions orchestrated by gRNA which evolved to give the virus a competitive advantage during assembly?

HIV particles assemble on the plasma membrane by incorporating Gag, Gag-Pol, Env and two copies of gRNA among other cofactors required for infectivity (Freed, 2015). The viral gRNA serves as a template for ribosomal translation of Gag and Gag-Pol while the Env is translated from a splice variant (Bolinger and Boris-Lawrie, 2009; Groom et al., 2009; Swanson and Malim, 2006). Binding of Gag to membrane, gRNA and tRNA (Bou-Nader et al., 2021; Göttlinger et al., 1989; Kutluay et al., 2014; Ono et al., 2004; Saad et al., 2006), loading of Gag-Pol within virions (Benner Bayleigh E. et al., n.d.; Hill M. K. et al., 2001), recruitment of gRNA and its dimerization (Chen et al., 2009; Chen Jianbo et al., 2016; Jouvenet et al., 2009; Keane et al., 2015; Nikolaitchik et al., 2021; Rein, 2019) and the choreography of HIV protease activation (Lee et al., 2012) has been extensively investigated. More importantly, the above studies have identified specific point mutations that abrogate each of the above interactions resulting in the release of non-infectious virions.

To further understand the budding mechanism of HIV, we created the conditions for competitive HIV budding by transfecting HEK293 cells with a constant amount of proviral DNA, fractionated between proviral DNA encoding the HIV gRNA (NL4.3) and proviral DNA encoding single point mutations abrogating specific protein-protein interactions. HIV gRNA is transcribed from NL4.3 proviral DNA and leads to assembly of infectious HIV virions in HEK293 cells. Our experiments probe how this gRNA competes with increasing quantities of the same gRNA incorporating specific mutants which have minimal effects on the gRNA structure and chemistry however abrogate the function of translated proteins in the same cell.

## Results

### Setting up the competitions

In our assay, the ratio of mutant/wildtype plasmids is varied from 0 to 100x and the released virions from HEK293 cells are assayed using western blots and their infectivity is measured in TZM-bl cells (Figure 1B). The 100x limit in dilution is dictated by the average number of proviral DNA in individual HEK293 cells that express proteins after transfection. Using a stochastic fluorescent approach (detailed in the supplementary methods) we have quantified the number of plasmids expressing proteins in individual HEK293 cells and show that each cell expresses proteins from an average of 100 plasmids with a distribution ranging from 20 to 300. Based on random incorporation of two gRNA’s within virions, the expected fraction of virions with gRNA homo dimer species can be calculated as shown with (light grey line in all figures) along with the gRNA hetero dimer (dark grey line). The equations governing these two theoretically predicted lines have been derived in the supplementary materials.

**Fig. 1.**
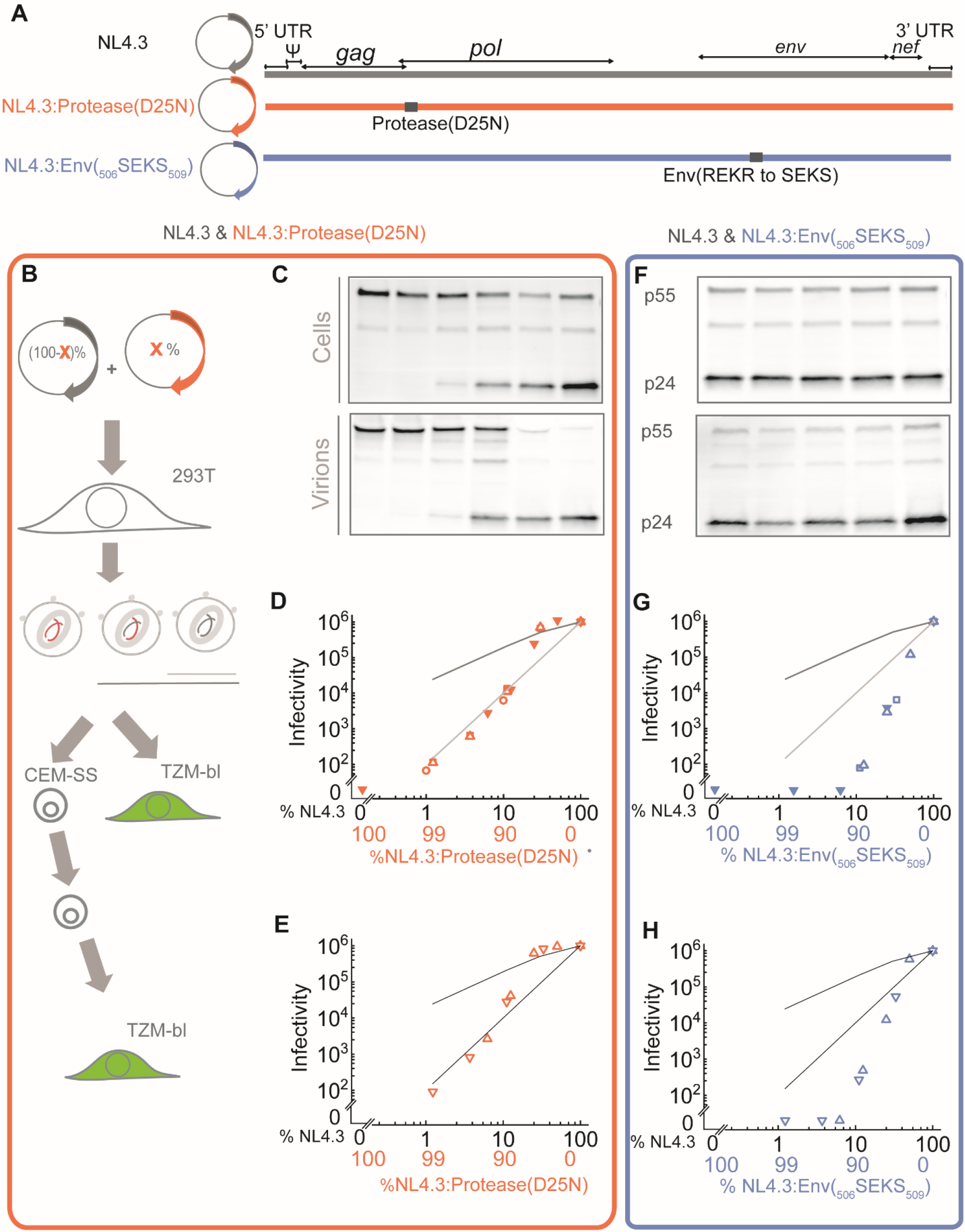
HIV assembly competitions between NL4.3 and NL4.3:Protease(D25N) and NL4.3:ENV(_506_SEKS_509_). The genome maps for NL4.3, NL4.3:Protease(D25N) and NL4.3:ENV(_506_SEKS_509_) (A). Schematic diagram of the competition experiments (B), Virions released by co-assembly of NL4.3 and NL4.3:Protease(D25N) in HEK293 cells assayed with western blots (C) and infectivity in TZM-bl cells (D) and infectivity after passage through CEM-SS cells (E) Virions released by co-assembly of NL4.3 and NL4.3:Env(_506_SEKS_509_) in HEK293 cells assayed with western blots (F) and infectivity in TZM-bl cells (G) and infectivity after passage through CEM-SS cells (H). Based on random incorporation of two gRNA’s within virions, the expected fraction of virions with gRNA homo dimer species can be calculated as shown with (light grey line in all figures) along with the gRNA hetero dimer (dark grey line). The equations governing these two theoretically predicted lines have been derived in the supplementary materials.

### Competitions with inactive Protease or fusion incompetent Env proteins

Since almost all treatments in ART include a protease inhibitor, the first competitions were set up to investigate the effect of an overwhelming quantity of protease mutants, specifically Protease(D25N) (Figure 1A). The 22kD HIV protease homodimer assembles from the 11kD protease precursor which is embedded within the Gag-Pol protein (Anderson et al., 2009; Pettit et al., 2005). Approximately 100 copies of Gag-Pol are embedded within individual HIV virions (Briggs and Kräusslich, 2011), the molecular mechanism of homodimer activation within virions is not well understood (Bendjennat and Saffarian, 2016). After activation however, the protease cleaves the immature HIV Gag lattice and results in formation of the mature HIV core which is essential for infectivity (Lee et al., 2012). The D25N mutation in HIV protease inhibits HIV protease activity and is one of the early mutations identified which resulted in release of noninfectious virions (Göttlinger et al., 1989). How many Gag-Pols are essential for activation of the protease is not known. In competitions shown in Figure 1, we simply asked how much excess NL4.3:Protease(D25N) is required to effectively ensure the suppression of infectivity in virions assembled from NL4.3 proviral DNA in HEK 293 cells.

In competitions between NL4.3 and NL4.3:Protease(D25N), the amount of released virions, as visualized in western blots, is constant regardless of the ratio of NL4.3 to NL4.3:Protease(D25N) (Figure 1.C). As the dilution of NL4.3 is increased, the infectivity of virions initially remains at or above ratios expected for virions packaging heterodimeric gRNA This trend continues up to 3X dilution of NL4.3 with NL4.3:Protease(D25N), at which point the infectivity starts dropping to the level expected from homodimer gRNA packaging. By 10X dilution of NL4.3 with NL4.3:Protease(D25N), the infectivity scales along the homodimer expectation and persists to the highest dilutions (100X) measured in our experiments. To verify the robust growth of the released virions, we also passaged the released virions in CEM-SS cells for 5 days at which point we co-cultured them with TZM-bl cells and measured the infectivity of the culture (Figure 1.E). These experiments confirmed that the virions released at high dilutions up to 100X were sufficiently fit and propagated within CEM-SS cells. The fact that virions assembling in cytosols with more than 100X excess of Protease(D25N) remain infectious is non-trivial.

We next tested Env. The ENV gene encodes the gp160 transmembrane protein which forms trimers after translation at the endoplasmic reticulum. Env(gp160) is proteolytically cleaved to form gp40 and gp120 which remain non-covalently associated within the trimer which gets packaged onto the budding HIV virions on the plasma membrane. The Env trimer is a target of neutralizing antibodies and is essential for virion fusion to the next host and infectivity. Env(_506_SEKS_509_) is deficient in proteolytic cleavage and virions incorporating this protein remain non-infectious (Pancera and Wyatt, 2005). We again asked how much excess of NL4.3: Env(_506_SEKS_509_) to NL4.3 proviral DNA can we add to effectively ensure that no infectious virions are assembled in HEK 293 cells (Figure 1).

Similarly to the competition between NL4.3 and NL4.3: Protease(D25N), the competition between NL4.3 and NL4.3:Env(_506_SEKS_509_) does not affect the virion release (Figure 1.F). As the dilution of NL4.3 and NL4.3:Env(_506_SEKS_509_) is increased, the infectivity of released virions drops and by 10X dilution is at the lowest limit detected in our experiments. In this regard, Env behave more like what we would expect for virions packaging from a pool of mixed proteins within the cell. It is important to note that both NL4.3:Env(_506_SEKS_509_) and NL4.3:Protease(D25N) transfected in HEK293 cells produce non-infectious virions. In competition with NL4.3, however, the infectivity data (Figure 1) shows curiously divergent behavior between NL4.3:Env(_506_SEKS_509_) and NL4.3:Protease(D25N). We therefore decided to investigate other non-infectious backbones.

### Competitions with Gag defective in membrane binding, recruitment of ESCRTs or gRNA defective in packaging and dimerization

One of the most severe mutants for HIV assembly is the NL4.3:Gag(G2A), which impairs the ability of Gag to bind membrane and results in loss of virion release as well as infectivity (Göttlinger et al., 1989). The data presented in Figure 2.B shows the infectivity of released virions in competitions between NL4.3 and NL4.3:Gag(G2A). As the dilution of NL4.3 with NL4.3:Gag(G2A) is increased, western blots show a reduction to release of virions (Figure 2.B). The infectivity of the supernatant also declines along the predicted line for gRNA homodimer species, suggesting that the only released virions with infectivity are virions which incorporate two NL4.3 gRNAs. The drop in infectivity along the homodimer gRNA line is unexpected, especially when compared to the complex behavior exhibited by NL4.3:Protease(D25N) (Figure 1D). Unlike the insensitivity of infectivity to low dilutions of NL4.3:Protease(D25N), the competition between NL4.3 and NL4.3:Gag(G2A) is sensitive from the lowest dilutions and follows the homodimer gRNA species line for two orders of magnitude in dilutions and three orders of magnitude in infectivity. At low dilutions, there is plenty of non-mutant Gag in the cytosol which can, in principle, bind and package all gRNAs. Based on extensive studies of the gRNA structure, the 2 nucleotide change introduced to create NL4.3:Gag(G2A) is not within the gRNA packaging signal therefore Gag molecules translated from NL4.3 gRNA should be capable to bind the gRNA from NL4.3:Gag(G2A) and package that with the same efficiency. This is clearly not supported by data shown in Figure 2.

**Fig. 2.**
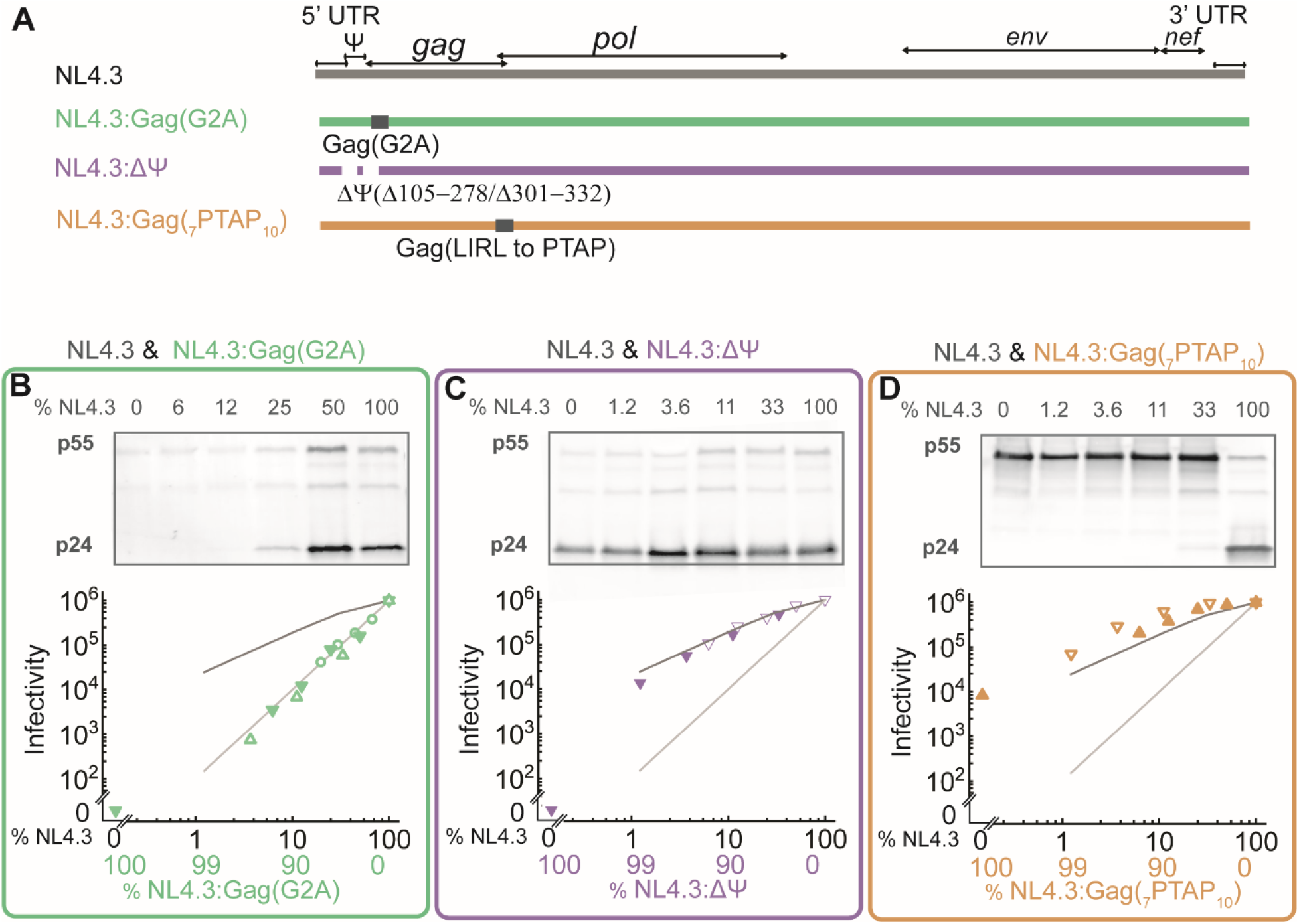
HIV assembly competitions between NL4.3 and NL4.3:Gag(G2A), NL4.3:ΔΨ(Δ105-278/Δ301-332) and NL4.3:Gag(_7_PTAP_10_). Genome maps (A). Virions released by co-assembly of NL4.3 and NL4.3:Gag(G2A) in HEK293 cells assayed with western blots and infectivity in TZM-bl cells (B). NL4.3 and NL4.3:ΔΨ(Δ105-278/Δ301-332) (C). NL4.3:Gag(_7_PTAP_10_) (D).

Given that endosomal sorting complexes required for transport (ESCRTs) catalyze the release of infectious HIV virions, Gag(_7_PTAP_10_) mutations abrogates the interaction of Gag with ESCRTs and results in release of mostly non-infectious virions(Huang et al., 1995). we next performed competitions between NL4.3 and NL4.3:Gag(_7_PTAP_10_) presented in Figure 2D. These were likely the least informative competitions within our study, since a large range of dilutions of the NL4.3:Gag(_7_PTAP_10_) could not effectively diminish the infectivity of the NL4.3. We therefore conclude that our method only works when NL4.3 is competed against mutants with very strong abrogation of infectivity, which is not the case for the PTAP mutation abrogating the interactions with ESCRTs.

HIV virions require incorporation of two gRNA molecules in order to be infectious. Deletion of sequences coding for RNA dimerization and the packaging signal NL4.3:ΔΨ(Δ105-278/Δ301-332) abolish viral infectivity, however they have a modest effect on Gag-gRNA interactions (Kutluay et al., 2014). Given that at high dilutions, many of our competitions behaved along the gRNA homodimer fraction, next we decided to compete the NL4.3 against NL4.3:ΔΨ(Δ105-278/Δ301-332) (Figure 2C). As expected the ratio of NL4.3 versus NL4.3:ΔΨ(Δ105-278/Δ301-332) proviral DNA has no effect on the amount of released virions, however as the dilution of NL4.3 with NL4.3:ΔΨ(Δ105-278/Δ301-332) is increased, the infectivity declines along the theoretical line expected from heterodimer gRNA incorporation suggesting that a NL4.3 derived gRNA can still dimerize and package along gRNA derived from NL4.3:ΔΨ(Δ105-278/Δ301-332). These results are slightly unexpected since they suggest that the dimerization signal is not required on both gRNA copies for dimerization to proceed which might suggest involvement of compensatory mechanisms.

### Developing models to better understand the infectivity data

The shape of the infectivity curves produced in our experiments (Figure 1 and 2) are distinct with clear features separated by orders of magnitude in both infectivity and dilution. The broader question we pose is how to interpret these infectivity graphs and what information can be obtained from these measurements. The HIV proteins competed in this study incorporate into HIV virions with varying stoichiometry. The Env proteins have the lowest stoichiometry within HIV virions ranging from 3-20 trimers interpreted from Cryotomography data (Brandenberg et al., 2015). Similarly, it is estimated that there are approximately 100 Gag-Pol proteins packaged in each virion, a number calculated based on the frameshifting ratio of Gag and Gag-Pol and the approximately 2000 Gag molecules estimated to be present in immature HIV virions based on CryoEM analysis (Briggs and Kräusslich, 2011). Cryotomography provides direct counting and stoichiometry measurements of proteins, approximately only 1-10% of released virions are infectious, therefore specifically measuring the protein stoichiometry of infectious virions remains challenging. Within each virion, there can be a mix of functional and non-functional proteins, how many functional proteins are required for infectivity also remains unanswered.

In our Monte Carlo model, we don’t model the spatial organization of the virions, trying to keep the models at bare minimum parameters. We treat each virion as a box, which needs to fill up with a set number of proteins from the cytosol. The box is made of different proteins, each with its own stoichiometry Env (10-30), Gag-Pol (80-120) and Gag (1500-2500). We did not assume an exact stoichiometry for either protein rather a range. Another parameter which was essential for our model was how many proteins from each kind needed to be functional for the box(virion) to be infectious. 10,000 boxes were randomly assembled for each stoichiometry. Fraction of boxes(virions) which had sufficient number of functional proteins was counted as infectious. Ratio of infectious to non-infectious virions was then calculated and plotted as infectivity data. In the trans packaging models, boxes are filled by randomly selecting proteins from the cell (Figure 3.A). The data shows that the slope of the infectivity decrease, is correlated with protein stoichiometry, with larger stoichiometry producing steeper infectivity drop off. The dilution at which the infectivity starts dropping is more sensitive to the fraction of proteins required for infectivity. This simple model with just two parameters therefore can capture the main behavior of the infectivity curves. There remained a problem however, because no parameter within the trans packaging model was capable of explaining the long infectivity tail observed in the NL4.3:Protease(D25N), NL4.3:ΔΨ(Δ105-278/Δ301-332) and NL4.3:Gag(G2A) infectivity presented in Figures 1&2. In (Figure 3.B&C) we show that a models where the gRNA packages some proteins in cis is required for explaining the long tail infectivity. Once some proteins arrive to the virions packaged in cis along the gRNA, one can either assume that these proteins are functionally equivalent to the rest of proteins that are packaged in trans (Figure 3B) or the cis packaged proteins are uniquely positioned and form the only functional proteins (Figure 3C). When trans and cis proteins are functionally equivalent, the behavior of the infectivity graphs becomes bi-phasic, where at low dilutions trans proteins are capable of rescue of all gRNA within the cytoplasm and creating infectious virions. At some larger dilutions however, when the majority of trans proteins within cytosol are dysfunctional, only functional cis packaged proteins are capable of rescuing the virion infectivity, which results in a direct correlation of released infectious virions with the fraction of packaged gRNA encoding functional proteins (Figure 3B). If only cis packaged proteins can rescue infectivity, the behavior of infectivity graphs become single phased with infectivity following the fraction of packaged gRNA encoding functional proteins from the lowest to highest dilutions.

**Fig. 3.**
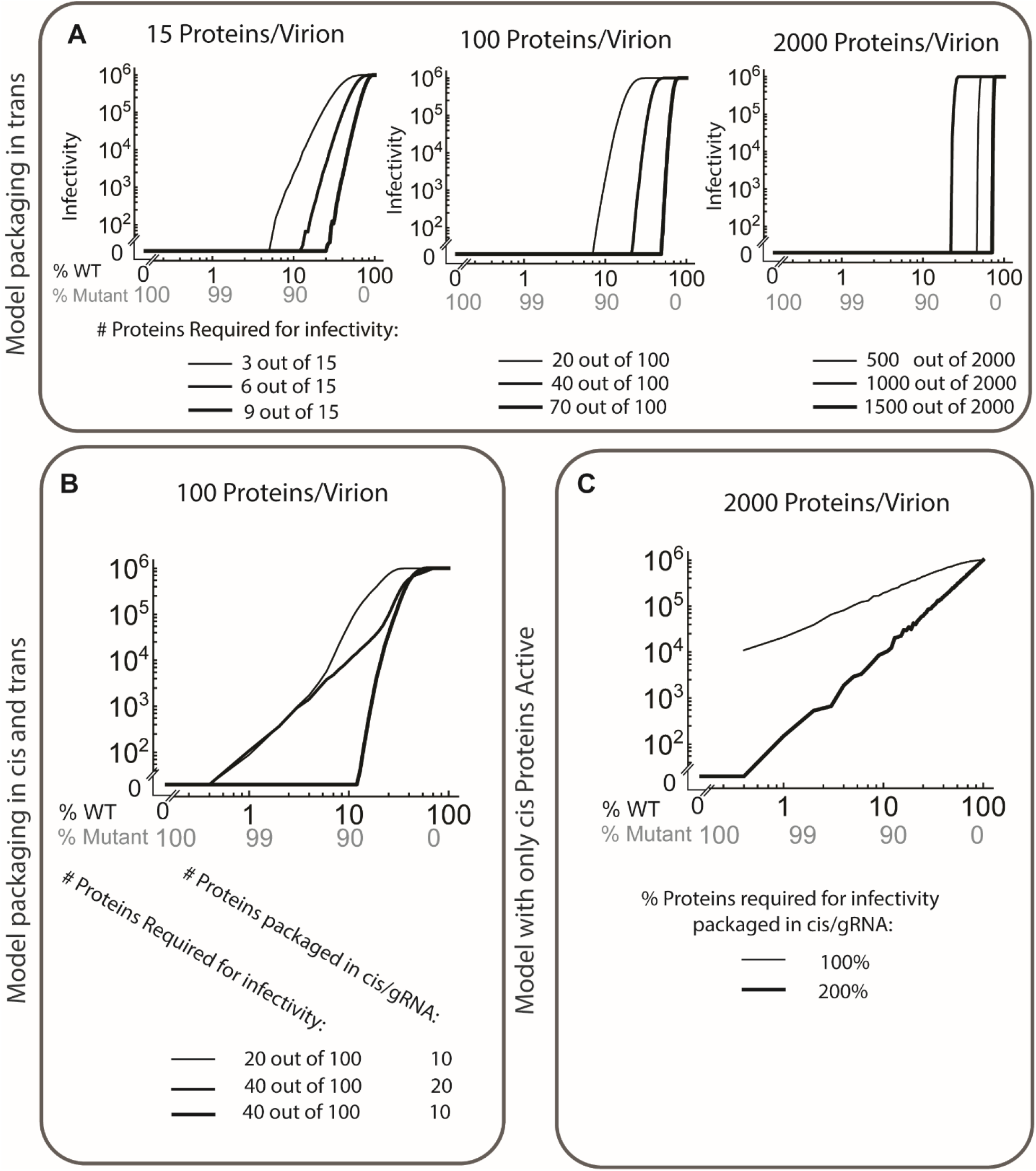
Monte Carlo models depicting various HIV packaging mechanisms. Models depicting packaging of a virion assuming only trans packaging of proteins along gRNA (A). Model assuming a fraction of proteins are cis packaged along gRNA of their origin (B). Predictions from a model where only cis packaged proteins can perform the required function essential for infectivity.

Using the predicted modeling curves shown in Figure 3, we specifically fitted the infectivity data from Figures 1 and 2 as shown along a model in Figure 4. In all simulations predict that each virion packages 100 Gag-Pols with at least 20 - 40 of them required for infectivity of the released virions, with each gRNA packaging 15 ± 5 Gag-Pol proteins in cis thus ensuring production of infectious virions from two WT gRNAs regardless of the trans packaging of the Gag-Pol on the membrane. Simulations also predict that approximately 70 Gag-Pols are packaged in trans, and these proteins can rescue the infectivity of other gRNAs that do not code functional Gag-Pols. The Monte Carlo simulations also demonstrate that the rescue in both G2A and ΔΨ is fully independent of trans packaging and only happens due to cis packaging of Gag or the inherent properties of the gRNA. Our findings illustrate that, in total HIV virions package in average 15 Env proteins, from which they need at least 9 of them to be functional for the virion to remain infectious. These measurements are broadly consistent with previous estimates for NL4.3 backbones assembling in HEK293 cells which found approximately 5 Env trimers are on each virion (Brandenberg et al., 2015).

**Fig. 4.**
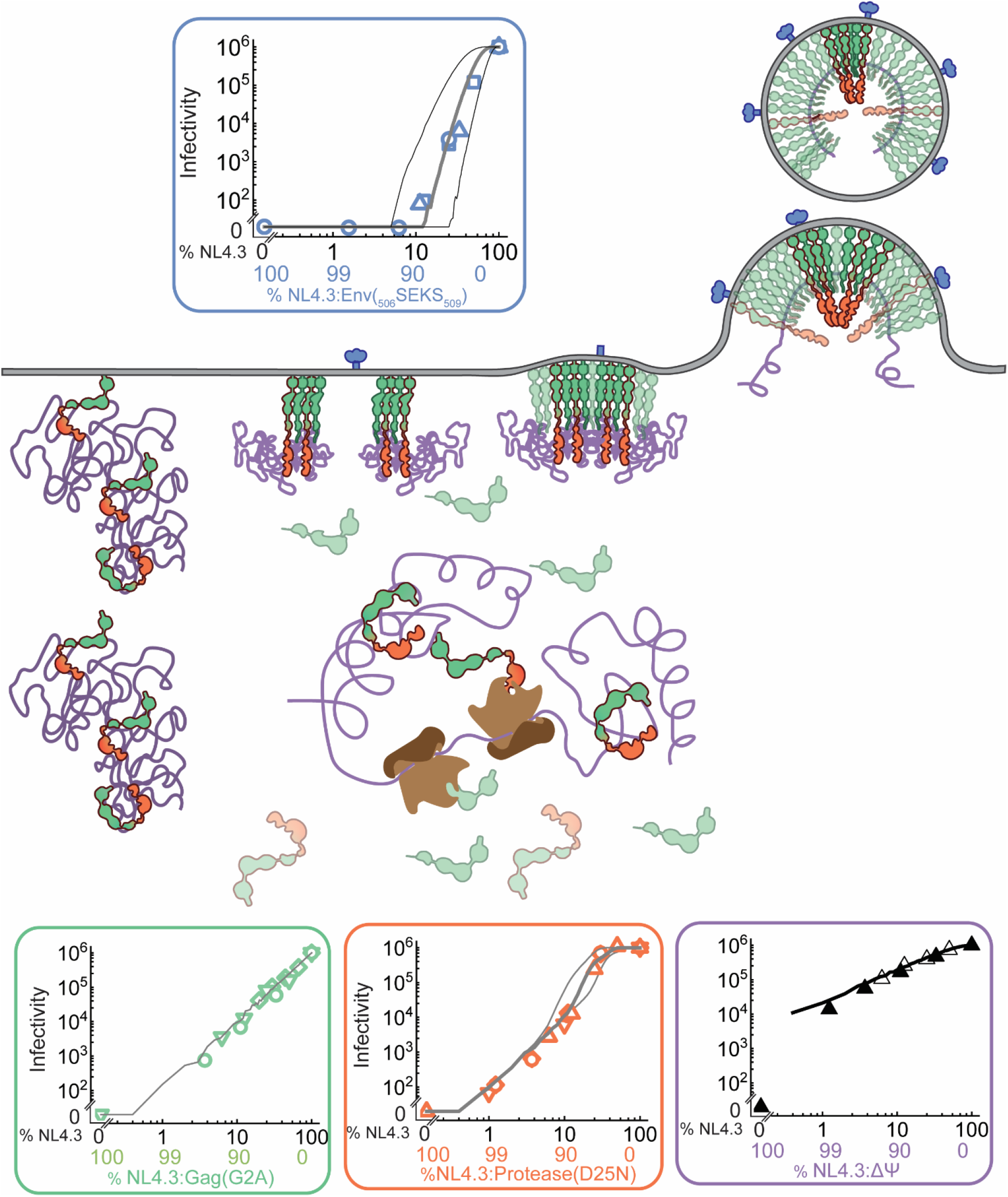
The model representation of the gRNA_1_:Gag-Pol_15_ complex. Inserts show infectivity data fitted by the corresponding Monte Carlo models. Env data is shown along models for trans packaging of 15 proteins with 6, 9 and 12 proteins essential for viral infectivity. G2A data is shown along Monte Carlo models with only cis packaging proteins capable of performing the function. Protease(D25N) data is shown along a model with a mix of cis and trans packaging. The lines represent 10, 15 and 20 proteins in cis with corresponding 20, 30 and 40 required for infectivity.

## Discussion

By quantifying the number of proviral DNA templates which can serve as viral initiation sites within individual HEK293 cells, we were able to setup competitions between various NL4.3 derived proviral DNA backbones and judge the competition based on the infectivity of released virions. Our method has a few advantages in that it focuses on the infectious virions and resolves some basic parameters such as stoichiometry of proteins, the number of functional proteins/virion required for infectivity and cis/trans packaging of proteins along gRNA. A side effect of our observation can be some new insight into the assembly initiation of infectious HIV virions. The initiation of assembly has been a complex and non-trivial issue (Bou-Nader et al., 2021; Göttlinger et al., 1989; Kutluay et al., 2014; Ono et al., 2004; Saad et al., 2006). One can propose based on our competition data in NL4.3:Gag(G2A) experiments and modeling which suggests that assembly initiation is driven only by cis packaged Gag-Gag-Pol, that cis packaged Gag-Gag-Pol within gRNA_1_: Gag-Pol_15_ complex initiate the assembly of virions at much lower cytosolic Gag concentrations, therefore ensuring specific packaging of HIV gRNA versus cytosolic mRNA. In this model the assembly is driven by gRNA and membrane templating which can in principle work at much lower cytosolic Gag concentrations. Its interesting to point out that our data is broadly consistent with previous reports showing some Gag-Pol packaging onto gRNA after translation(Benner Bayleigh E. et al., n.d.).

We also list potential caveats below, however none of the caveats significantly impact our study and in principle, this method can therefore, find its application in studying other enveloped RNA viruses or other essential viral enzymes/structures within HIV. In this regard, once a significant mutation resulting in full inactivation of the virions is identified, the competition between that mutant and the WT virions will result in better understanding of the number of proteins incorporated within virions and the mechanism of packaging cis/trans.

Caveats: It’s important to note that while HEK 293 cells provide a convenient cytosol for competitions between up to 100 proviral DNA units, in these competitions, some of the essential transcription machinery required for optimum packaging of HIV virions maybe saturated. While we do not expect that this saturation had a significant effect on our results, it is likely that the NL4.3 gRNA maybe even more competitive if operating within a dynamic range where all transcriptional elements are in abundance (Singh Gatikrushna et al., 2022). The other important issue is that HEK293 cells are not natural hosts of HIV and therefore our method is blind to further evolutionary aspects of gRNA which may enable even more efficient escape from the primary cells.

Outlook: We present a method which has a distinct capability in resolving a few very important aspects of infectious HIV assembly. It’s important however to highlight, that our method, is not capable of answering mechanistic questions as to how the gRNA_1_: Gag-Pol_15_ is stabilized within the cytosol, how long it remains stable and what are the molecular interactions which keep the complex together. We also do not resolve the number of Gag molecules associated with the gRNA_1_: Gag-Pol_15_ complex although we assume that there must be at least some. The questions of complex stability are even more profound, when one realizes that the complex must remain together while multiple ribosomes read and translate the *gag, gag-pol* region likely displacing any specific binding interactions. Understanding these molecular interactions will take development of further imaging modalities, methods and possibly new ways of thinking about HIV.

## Acknowledgments

The authors acknowledge fruitful discussions with Dr Michael Vershinin during conception of the idea and writing of the manuscripts. Dr Emanuele Cocucci and Dr Till Böcking for fruitful discussions. Dr Janet Iwasa generously shared models which we edited to create figure 4. The p24 antibodies used in western blots, NL4.3 backbone, TZM-bl and CEM-SS cells were obtained from the NIH HIV reagent program. This study was supported by NIH R01 AI150474 to (SS).

## Author contributions

Conceptualization: SS, IS

Methodology: SS, HD, BP

Investigation: HD, BP, BM, AP, WP

Funding acquisition: SS

Project administration: HD, SS

Supervision: SS

Writing – SS

Writing – SS, HD, BP and IS

## Data and materials availability

“All data are available in the main text or the supplementary materials.”

## Supplementary Materials

Materials and Methods

Supplementary Text

Figs. S1 to S3

